# Roles of linker region flanked by transmembrane and peptidoglycan binding region of PomB in energy conversion of the *Vibrio* flagellar motor

**DOI:** 10.1101/2023.11.13.566875

**Authors:** Yusuke Miyamura, Tatsuro Nishikino, Hiroaki Koiwa, Michio Homma, Seiji Kojima

## Abstract

The energy converting complex of the sodium-driven flagellar motor in bacteria comprises two proteins, PomA and PomB, whose transmembrane regions form ion conducting channels and is called the stator complex. The transmembrane protein PomB is attached to the cell wall by its periplasmic region and has a plug segment following the transmembrane helix to prevent ion flux. PomB (Δ41-120), which lacks the periplasmic region from E41 to K120 immediately following its transmembrane region shows similar motility as that of wild-type PomB. In this study, three deletion mutants after the plug region, PomB (Δ61-120), PomB (Δ61-140), and PomB (Δ71-150), were generated and those deletion mutants were examined for their functionality. PomB (Δ61-120) conferred similar motility as that of the wild-type protein, whereas the other two mutants showed almost no motility in soft agar plate; however, we observed some swimming cells with speed similar to that of the wild-type cells. To observe dominance of wild-type proteins, we introduced the two PomB mutants into a wild-type strain, and its ability to swim was not affected by the mutants. Then, we purified the mutant PomAB complexes to confirm the stator formation. When we introduced the PomB mutations in the plug region, the reduced motility by the deletion was rescued, suggesting that the stator was activated. Our results indicate that the deletion prevents stator from transformation to an active form; however, the linker and plug regions from E41 to S150 are not essential for the motor function of PomB but are important for its regulation.

**IMPORTANCE:** The stator complex of flagella consists of PomA and PomB proteins and interacts with the rotor complex. PomB has a peptidoglycan binding (PGB) domain to fix the stator for generation of torque. PomB is attached to the cell wall only when the stator is activated by interaction between the cytoplasmic region of PomA and the rotor protein FliG. The flexible linker of PomB, which is a naturally unfolded region, is flanked by the peptidoglycan-binding (PGB) domain and transmembrane region. The plug region, which interacts with the periplasmic loops of PomA to prevent activation of the stator, is located next to its transmembrane region. In this study, we reveal the role of the flexible linker in activation of the stator complex.

## INTRODUCTION

Many bacteria require motility for survival. Some of them have flagella, which are fibrous motile organelles on the cell surface and are responsible for movement. Marine bacterium *Vibrio alginolyticus* used in this study or *V. parahaemolyticus*, a close phylogenetic relative, have a single polar flagellum and multiple peripheral lateral peritrichous flagella (1-3). The polar flagellum is rotated by sodium ion flux to swim in sea water. Lateral flagella are rotated by proton flux to swarm on surface of fish. At the base of the flagellum, a flagellar motor serves as an engine. The motor consists of the stator complex, which is an ion energy conversion unit, and a rotor that generates torque through interaction with the stator complex (4). The stator complex is a membrane protein complex comprising a PomB dimer surrounded by a pentamer of four transmembrane (TM) protein PomA (5). The cytoplasmic region of PomA interacts with the rotor protein FliG to generate rotational forces (6).

The three-dimensional structures of the proteins in stator complex have been recently reported and a stator rotation model was proposed, in which the stator’s own clockwise (CW) rotation conjugated with the ion current is transmitted to the rotor via a gear consisting of conserved charged residues that causes rotation of the rotor ring (7, 8). Existence of this model has not been directly demonstrated; however, we have reported that a helix located immediately after the transmembrane region of PomB, acting as a plug to interact with the stator channel, may mechanically inhibit the rotation of the PomA ring (9). Initially, this plug region was considered to be the lid of the ion channel of the stator complex (10, 11). Overexpression of the stator complex proteins does not alter the growth of the cell; however, cells over-expressing mutant stator protein lacking the plug region are unable to grow. The over-expression of PomAB and MotAB complex lacking the plug region in *Escherichia coli* affected Na^+^ and H^+^ flux as shown by increased Na^+^ concentration, which was measured using atomic absorption spectroscopy (12), and H^+^ concentration, which was measured using a pH indicator protein pHluorin (13). In contrast, absence of linker region does not affect the rotational mechanism of the flagellar motor; however, extensive structural change of the linker region is required for regulating the function of the stator complex or for its proper positioning around the rotor via the peptidoglycan-binding region (PGB domain) of the PomB extracellular region (14). The structures of PGB domains and cytoplasmic and transmembrane regions of proteins in the stator complex have been solved (7, 8, 14-17); however, that of the linker region has not been solved probably because of its flexibility given that it is an intrinsically disordered region. In this study, we focused on elucidating structure and function of the linker region.

## RESULTS

### PomB linker deletion mutants and their phenotypes

Three mutant PomB proteins, PomB (Δ61-120), PomB (Δ61-140), and PomB (Δ71-150), were generated by deleting their linker regions (Fig. 1A and Fig. S1). These deletion PomB constructs were expressed together with wild-type PomA from the arabinose-inducible plasmids in the *pomAB* deletion strain NMB191 and their motility in soft agar plate was examined. The same constructs were introduced into *E. coli* DH5α cells and the bacterial growth was observed by the arabinose induction in LB broth containing 3% (w/v) NaCl. Strains with PomB (Δ61-140) and PomB (Δ71-150) did not exhibit the swimming ring; however, the strain with PomB (Δ61-120) exhibited the swimming ring similar to the strain producing wild-type PomB (Fig. 1B). When cells cultured in liquid medium were observed using dark-field microscopy, approximately 10% of the total bacterial cells producing PomB (Δ61-140) and PomB (Δ71-150) were found to be motile and their swimming speeds were similar to those of the wild-type cells (Fig 2).

**FIG 1.**
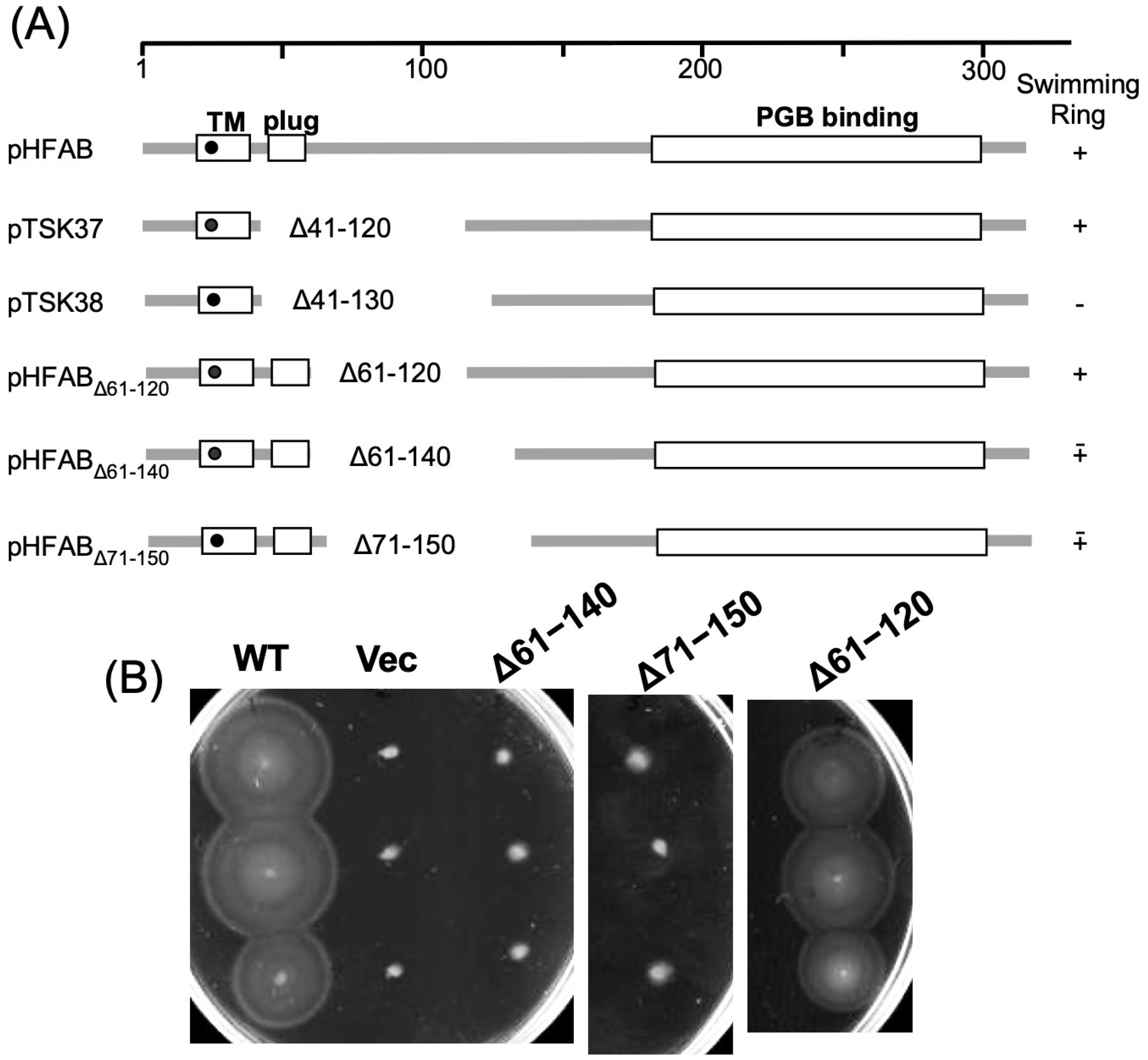
(A) Schematic of the primary structure of PomB fragments. The numbers under the line indicate the amino acid number and the black filled circle in the transmembrane (TM) region indicates Asp residue which is a sodium binding site. (B) Motility of the PomB deletion mutants on soft agar plates. The *pomAB* deletion mutants (NMB196) containing pHFAB (WT), pBAD33 (Vec), pHFAB_Δ61-120_ (Δ61-120), pHFAB_Δ61-140_ (Δ61-140), and pHFAB_Δ71-150_ (Δ71-150) were inoculated on soft agar plates at 30 °C for 5 hours.

**FIG 2.**
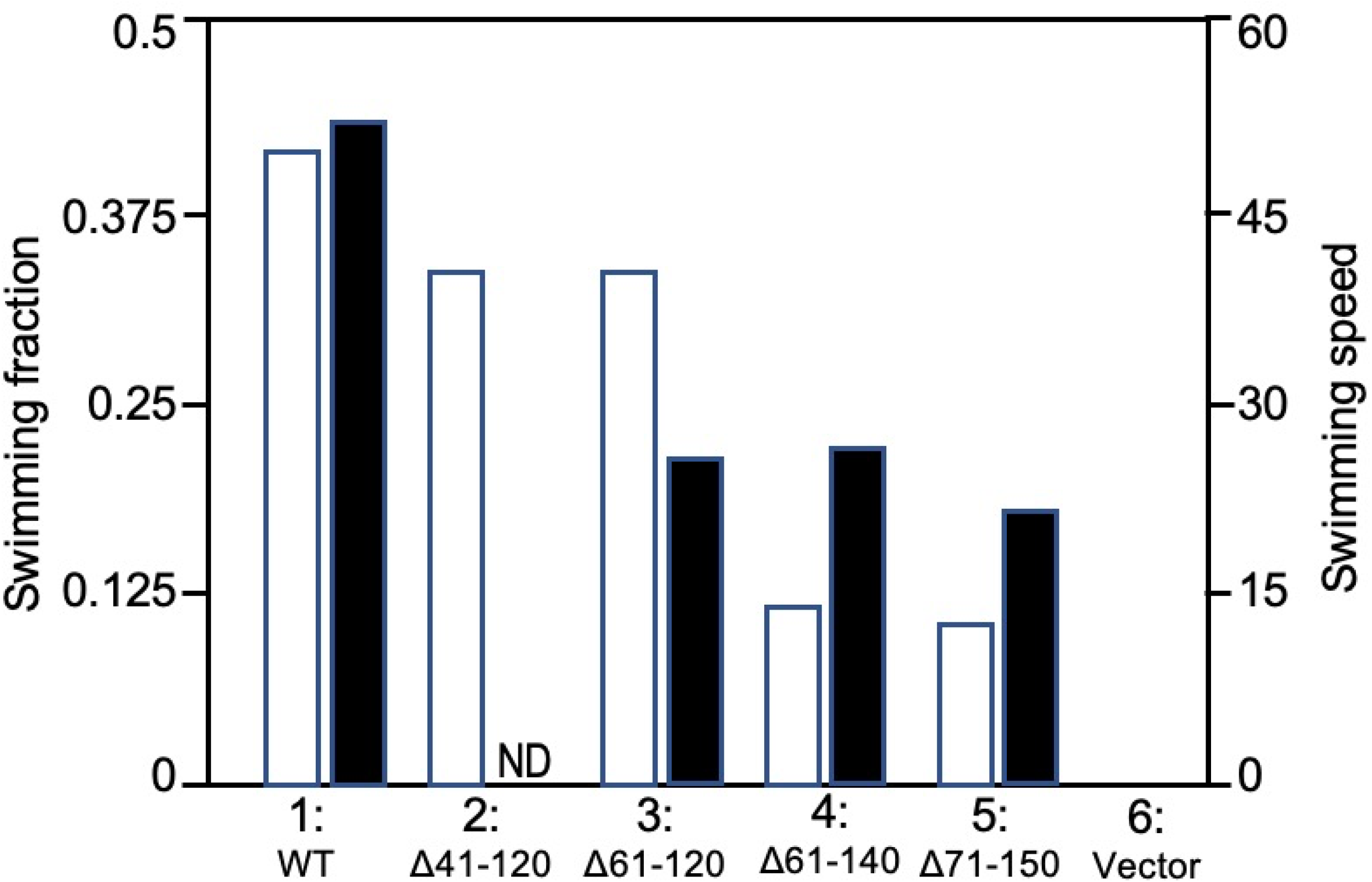
Swimming profiles of PomB deletion mutants. *pomAB* deletion mutant (NMB196) cells containing pHFAB (WT: 1), pTSK37(Δ41-120: 2), pHFAB_Δ61-120_ (Δ61-120: 3), pHFAB_Δ61-140_ (Δ61-140: 4), pHFAB_Δ71-150_ (Δ71-150: 5), pBAD33 (Vector: 6) were grown in the VPG medium at 30 °C up to log-phase and their swimming profiles were observed using dark-field microscopy. Open and closed bars show the swimming fraction (ratio) and swimming speed (μm/sec), respectively. ND; not tested. In cells containing pBAD33: 6, no motile cells were detected.

Dominance of the wild type proteins was verified by expression of mutant PomB in the *Vibrio alginolyticus* strain VIO5, which is the wild-type strain for the motility of polar flagella. We showed that motility in soft agar plate was not affected by expression of the mutant PomB proteins (Fig. S2A). The plug-deficient PomB Δ41-120 caused inhibition of growth as reported previously (12), whereas the mutant strains generated in this study did not show growth inhibition (Fig. S2B). Furthermore, we confirmed the production of the mutant PomB proteins in the cells by immunoblot, and both PomA and PomB mutant proteins were detected as the wild-type PomA and PomB (Fig. S3). The molecular weights estimated using SDS-PAGE are correspond to the mutant proteins predicted based on amino acid sequence deduced from DNA sequence.

### The stator complex comprising mutant PomB and wild-type PomA

We examined properties of the PomAB complex comprising mutant PomB. We established the purification procedure of the stator complex expressed in *E. coli* using a cold expression vector (18). PomB deletions were introduced into the vector, pColdIV-pomAB-his_6_ which contained the *pomA* and *pomB* genes and was renamed as pCABH11 in this study.

Stator complexes composed of wild-type PomA and mutant PomB (Δ61-120), PomB (Δ61-140), or PomB (Δ71-150) were isolated. Their purification profiles were similar to those of the wild-type PomAB complex (Fig. S4). When the stator complex was purified using size-exclusion chromatography, the complex was eluted at the volume corresponding to a molecular weight of about 430 kDa of molecular weight (Fig 3). This is much higher than the estimated molecular weights of five PomA and two PomB (Δ71-150) subunits, which is estimated to be 189 kDa. This difference is probably owing to association of complexes with detergent. We observed the complex using electron microscopy by negative staining. It is difficult to distinguish the structural differences between the stator complexes composed of the wild-type and the mutant PomAB and we could detect similar particles in both complexes (Fig. S5). The stator complexes consisting of PomA and PomB (Δ41-120) could not be purified with this method, probably because PomA dissociated easily from the complex when the plug region of PomB is deleted (19). At least, the three deletion mutants of PomB made in this study formed stable stator complexes with wild-type PomA.

**FIG 3.**
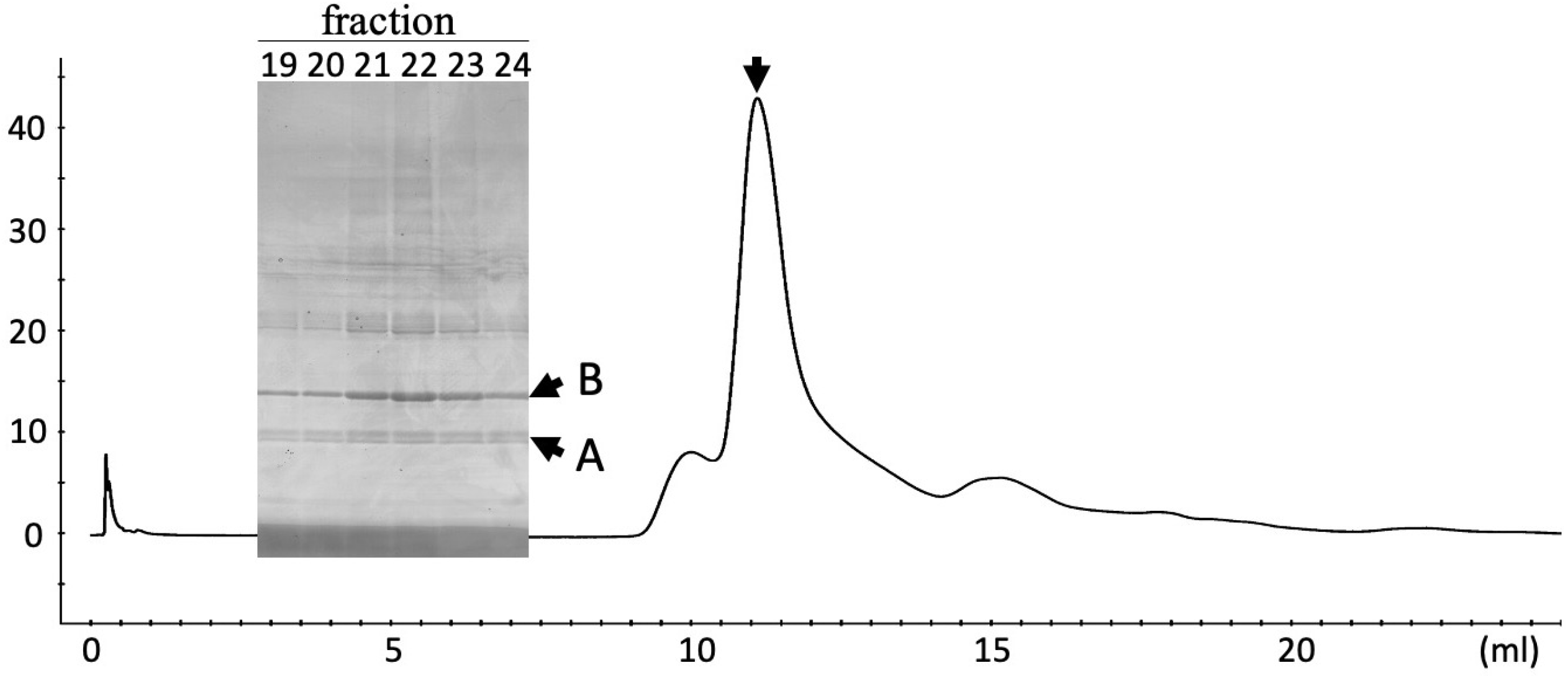
Elution profile of gel filtration chromatography of the PomAB complex. *E. coli* cells containing pCABH14 were used to produce PomA and PomB (Δ71-150) deletion mutants which were purified using Ni-affinity purification using a His-tag for PomB. Protein sample (0.5 mL) were injected into the size exclusion column (BIO-RAD ENrichTM SEC650) and eluted using the 20TN100 buffer containing 0.0025% LMNG at a flow rate of 0.5 mL/min. Elution profiles were monitored based on the absorbance at 280 nm. The peak position indicated by the arrow is 11.1 mL. Proteins from fractions 19-24 of the elution were separated using SDS-PAGE, followed by staining with Coomassie brilliant blue. A and B indicate the protein bands of PomA and PomB, respectively.

### Suppressor mutation for recovery of motility in the linker mutants

We isolated mutants with the ability to form swimming rings on soft agar from the *V. alginolyticus pomAB* null mutant strain (NMB191) expressing PomB (Δ71-150) or PomB (Δ61-140) with wild-type PomA. Four mutants showing swimming rings were isolated from PomB (Δ71-150) and from PomB (Δ61-140) expressing strain each. When recombinant plasmids extracted from the mutants were used for re-transforming the parent strain of NMB191, the two mutants (number 3 and 4) showed a swimming pattern similar to that of the source mutant cells and other mutants did not recover their swimming ability (Fig. 4). Recombinant plasmids from mutants 3 and 4 were sequenced to examine the sequence at mutation sites. We identified the M20I mutation of PomA in the recombinant plasmid from mutant 4; however, mutant 3 contained an unfamiliar sequence.

**FIG 4.**
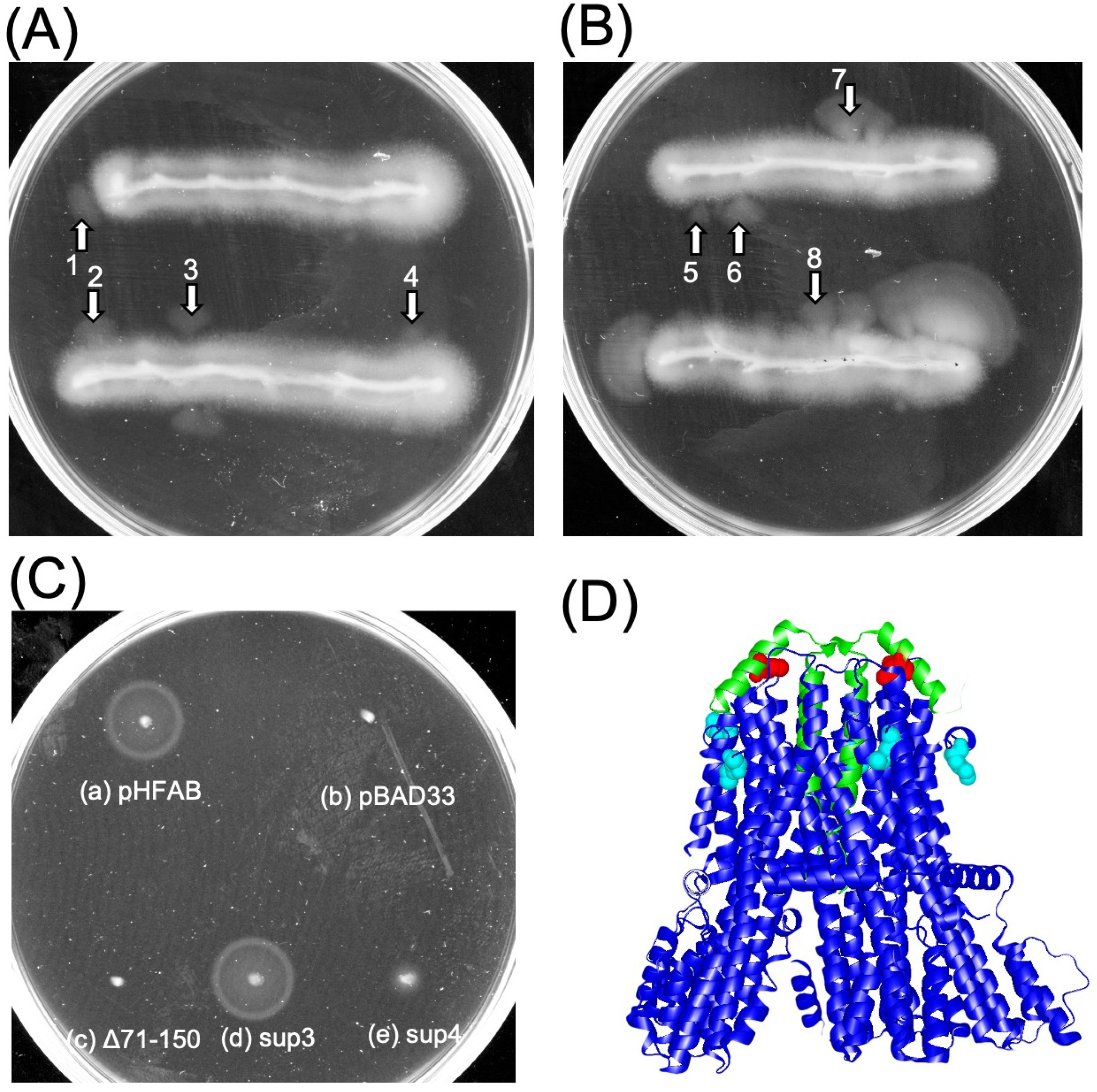
Isolation of the suppressor mutants. The *pomAB* cells containing pHFAB_Δ71-150_ (Δ71-150) (A) and pHFAB_Δ61-140_ (Δ61-140) (B) were streaked on the VPG soft agar plates and incubated at 30 ℃ overnight. Arrows indicate isolated mutants from the plates. (C) *pomAB* cells transformed using pHFAB (a), pBAD33 (b), pHFAB_Δ71-150_ (c), recombinant plasmids isolated from mutant 3 (sup3: d) of (A) or from mutant 4 (sup4: e) of (A). Colonies were inoculated on soft agar plates and incubated at 30 ℃ for 4 h. (D) Position of the suppressor mutation. The I50 residue of PomB and the M20 residue of PomA are shown by space-filling with red and light blue respectively, in a ribbon model of *V. alginolyticus* PomA/B complex (PDB: 8BRI) in which PomA and PomB are shown in blue and green, respectively.

Next, we introduced additional mutations, PomB-I50E and PomB-I50C, into the PomB (Δ71-150) mutant and examined the motility of mutant cells (Fig. 5). The *pomAB* mutant cells producing wild-type PomB and PomB (Δ71-150) with PomA were motile in ca. 44% and 13% of examined populations, respectively. In contrast, PomB-I50E and PomB-I50C rescued the motility of PomB (Δ71-150) in ca. 32% and 39% of examined bacterial cells, respectively. From these results, it can be inferred that PomB-I50E and PomB-I50C mutations assisted activation of the stator complex.

**FIG 5.**
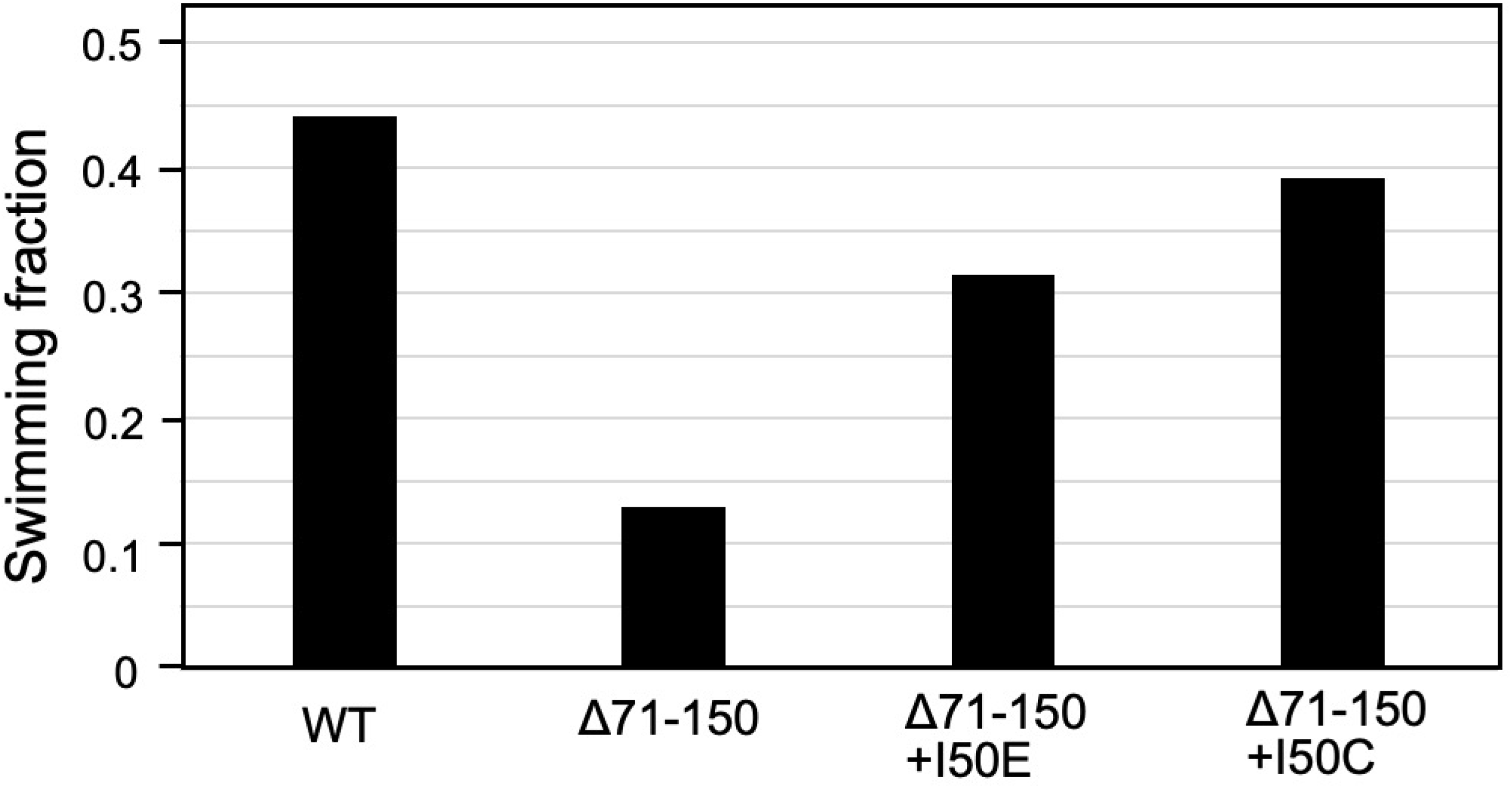
Suppressor phenotype caused by PomA mutations in PomB deletion mutants. Cells containing pHFAB (WT) or pHFAB_Δ71-150_ (Δ71-150) and with the I50E or I50C mutation of PomA were grown in the VPG medium at 30 ℃ up to log-phase and their swimming profiles were observed using dark-field microscopy.

## DISCUSSION

The flagellar motor of bacterial cells is a biological nanomachine for rotation of the filament extended from the cell surface, which confers motility to the cells. The motor has an energy converting stator complex composed of membrane proteins, PomA and PomB in *Vibrio* cells and is attached to the peptidoglycan layer in the periplasmic space by the periplasmic region or PGB domain of PomB (3). The PGB domain is connected by a linker, which is composed of approximately 130 amino acids, to the membrane-bound structure of the stator complex (10). PomB is attached only when the stator complex is activated by interaction between the cytoplasmic region of PomA and the rotor protein FliG (20, 21). It has been shown in marine *Vibrio* species that the deletion of 80 residues in PomB, from E41 to K120 (Δ41-120) of the linker and plug region between the transmembrane region and the PGB domain, retained motility, whereas bacteria with the PomB (Δ41−130) deletion were not motile (10). We speculated at the time that the sequence from 120 to 130 may be important for the motor function. The PomB plug region is located immediately after the TM region from V44 to A57 and forms an α helix. The conservative Serine 40 residue (S40) is placed at the boundary of TM region to break the α helix of TM. In this study, we showed that the motor function of the stator complex was retained even when additional 20 residues (131-150) were deleted in the presence of the plug region as PomB (Δ71-150). Thus, these regions do not appear to have an essential function.

PomB (Δ71-150) and PomB (Δ61-140) mutants showed almost no swimming ring formation on soft agar plates; however, we observed more than 10% motile cells using dark-field microscopy. The speed of motile cells was almost the same as that shown by cells with wild-type PomB. This suggests that the stator complex deletion mutant has potential ability to drive the motor; however, most of these complexes are not in the active state. First, we isolated the revertant mutants. It was simple to obtain mutants that exhibited swimming rings larger than those of the parent cells (Fig. 4AB). When we isolated recombinant plasmids from the revertant mutants and retransformed the *pomAB* mutant cells, most plasmids isolated from the revertant cells could not rescue motility. This may mean that the mutations were present in the chromosome of the *pomAB* mutant. We could obtain two recombinant plasmids isolated from the revertant cells that conferred motility to the *pomAB* mutant and the mutation site of one was altered from Met20 of PomA to Ile. This site has been identified in the structure of the PomAB complex (Fig. 4C) very recently (17). The position of PomA-M20 is located at the end of the TM1 helix and corresponds to the outer surface of the lipid bilayer. This helix appears to be flexible in the stator complex as suggested by the structure of PomAB reconstituted in lipid nanodiscs (17). We speculated that this mutation might affect the plug interaction between PomA and PomB. We have shown that the plug region of PomB terminates the ion influx by blocking the rotation of the rotor similar to a spanner (9).

We have shown that cells producing the PomAB complex with the mutations F47, I50, or M54 in the plug region show defective growth. Cells with mutations at I50 of PomB and M169 of PomA, which is in the periplasmic loop between TM3 and TM4, appear to interact or to be very close to each other (9). The residues (I50, M54 and F58) from the plug motif are deeply inserted into this cleft to establish hydrophobic interactions, and that the PomB F47 aromatic ring is sandwiched between the pyrrolidine ring of P172 and the side chain of M169 from PomA, via CH-π interactions, further stabilizing the plug motif (17). When we introduced the I50C or the I50E mutation in PomB (Δ71-150), the motility of cells was recovered (Fig. 5). This is probably because the interaction between the plug and loop of TM3 and TM4 was released and activated. Structural analysis has shown that the flexibility of the TM1 helices in the inactive stator unit is probably intrinsic, which might be functionally important during stator unit activation (17). The evidence shown in this study supports our hypothesis.

The linker region is important for interaction with FliL, which is not an essential protein for the motor function but is necessary for functioning of the *Vibrio* polar flagellar motor in highly viscous media (22, 23). The *fliL* mutant of *Helicobacter pylori* lost motility. *Helicobacter* FliL sequence is very different from that of *Vibrio* FliL; however, their protein structures are very similar (24). The linker region and the plug region of PomB are surrounded by a decamer of FliL molecules, which form a ring structure (25). It was shown in *H. pylori* that the FliL ring structure was coaxially sandwiched between the MotA ring and dimeric periplasmic MotB region, and there was the density with the central hole of the FliL ring and with the plug/linker region of MotB as the extended form of active conformation (24). We speculate that the stator complex in the deletion mutants used in this study might not be surrounded by FliL proteins.

Some reports suggested that FliL interacts with MotA, MotB, and FliF, though the interaction sites of FliL remain unknown (26, 27). In the structure of the PomAB stator complex determined using cryo-EM, the residues beyond Q61 were not visible (17). In the crystal structure of the C-terminal periplasmic region of PomB determined using X-ray crystallography, the residues up to M153 were not visible even when the fragments from D121 and T135 of PomB were crystalized (14). We have shown that the C-terminal fragment of *S. enterica* MotB from A78 contains highly mobile region composed of approximately 40 amino acid residues using 1H-15N Hetero-nuclear Single Quantum Coherence (HSQC) spectroscopy (15). As expected, the region from Q61 to M153 of PomB is highly mobile and it is difficult to determine its structure. As for the structure of *H. pylori* MotB, several truncated fragments were crystalized and one of them revealed the structure of the linker region of MotB (28) with a small secondary structure. The structural prediction by AlphaFold2 for the PomB fragment from 1 to 150 indicated that there is no secondary structure in the linker region of PomA except for the C-terminal region of the fragment (Fig. S6). The region from Q61 to M153 of PomB seems to be the intrinsically disordered region and is involved in the regulation of motor function; however, it is not essential for generation of torque in the motor.

## MATERIALS AND METHODS

### Bacterial strains and plasmids

Bacterial strains and plasmids used in this study are shown in Table S1. *E. coli* was grown in LB [1% (w/v) bactotryptone, 0.5% (w/v) yeast extract, and 0.5% (w/v) NaCl]. For growth on a solid medium, 1.25% (w/v) agar was added to LB. If necessary, chloramphenicol and ampicillin were added to a final concentration of 25 μg/ml and 100 μg/ml, respectively.

*V. alginolyticus* was cultured at 30°C in the VC broth [0.5% (w/v) hipolypeptone, 0.5% (w/v) yeast extract, 0.4% (w/v) K_2_HPO_4_, 3% (w/v) NaCl, and 0.2% (w/v) glucose] or VPG medium [1% (w/v) hipolypeptone, 0.4% (w/v) K_2_HPO_4_, 3% (w/v) NaCl, and 0.5% (w/v) glycerol]. For growth on solid medium, 1.25% (w/v) agar was added to VC broth. A VPG soft agar medium was prepared by adding 0.25% bactoagar, respectively. If necessary, chloramphenicol was added to a final concentration of 2.5 µg ml^−1^, or arabinose was added to final concentrations of 0.02 or 0.2% (w/v) for producing proteins.

### DNA manipulation and mutagenesis of *pomB*

Routine DNA manipulation was performed according to standard procedures. Deletion mutations in *pomB* were introduced into plasmids using a one-step PCR-based method as previously described (10). Each mutation was verified using DNA sequencing.

### Motility assay

One microliter of overnight cultures of *V. alginolyticus* NMB191, containing recombinant plasmids at 30°C in the VC medium was spotted onto VPG soft-agar plates [VPG medium and 0.25% (w/v) bactoagar] containing 0.02% (w/v) arabinose. These plates were incubated at 30°C for 5 h.

For analysis of bacterial swimming, these overnight cultures were diluted 100-fold in fresh VPG medium with 0.02% or 0.2% (w/v) arabinose. After incubation at 30°C for 4 h, cultured cells were diluted 100-fold in the VPG medium and were observed using dark-field microscopy. Movies of cell motility were recorded and the swimming fractions and speeds of the cells were measured using software for motion analysis (Move-tr/2D, Library Co.) or using motion analysis plug-in of image J.

### Purification of the PomAB complex

*E. coli* harboring the PomAB expression vector pCABH11 (pCold4-pomApomB-His6) and the deletion derivatives were cultured overnight in LB at 37°C. After 100-fold dilution in fresh LB, the fresh culture was incubated at 37°C for 2.5-3.0 h until the OD_660_ reached 0.4-0.6 and then isopropyl β-D-1-thiogalactopyranoside (IPTG) was added at a final concentration of 0.5 mM. This culture was transferred to ice-water for 30 min and was incubated at 16°C for 20 h. Cells were collected by centrifugation (4,600 × *g*, 10 min) and resuspended in 30 ml 20TK200 buffer (20 mM Tris-HCl [pH 8.0], 200 mM KCl) containing 1 mM ethylenediaminetetraacetic acid (EDTA). Cells were lysed using the French press (9,000 – 10,000 psi, OTAKE Works) or by sonication (large probe, power = 6, 60 s, 3 times, duty cycle 50%). Cells that were not lysed were removed by centrifugation at 10,000 × *g* for 10 min at 4°C, and were resuspended in 20 mL 20TK200 buffer containing 1 mM EDTA, and sonication was repeated.

Resulting supernatants were collected, MgCl_2_ and imidazole were added at a final concentration of 2 mM and 20 mM, respectively, cells were then centrifuged at 12,000 × *g* for 10 min at 4°C, and the supernatant was further ultracentrifuged at 154,000 × *g* for 60 min at 4°C. The supernatant and pellet are defined as the soluble cytoplasmic fraction and the membrane fraction, respectively. The precipitate of the membrane fraction was resuspended in 20 mL 20TK200 and was shaken in a cold room overnight. 2 mL 10% LMNG solution and 100 µL 2M imidazole was added and centrifuged at 10,000 × *g* for 10 min. The supernatant was used for Ni-affinity purification using a His-tag.

Protein samples were loaded onto a HiTrap TALON column (5 mL, GE Healthcare) connected to an AKTA prime system (GE Healthcare) and washed with the 20TK200 buffer or 20TN200 buffer containing 0.01% LMNG and 10 mM imidazole. The protein was eluted and collected using a 10 mM to 300 mM imidazole linear gradient. Peak fractions were concentrated by centrifugation using the 100K Amicon filter device (Millipore).

Protein samples were centrifuged (150,000 × g for 10 min at 4°C) and loaded onto an Enrich SEC 650 10 × 300 column (Bio-Rad) using the AKTA explore system (GE Healthcare) at a flow rate of 0.5 ml/min. Eluted samples were verified using SDS-PAGE and peak fractions were concentrated as described before without the addition of 2-mercaptoethanol.

### Detection of PomA and PomB protein from whole cell lysate

Whole cell lysate of the *Vibrio* cells was prepared by adding an SDS-PAGE loading solution to cell suspension whose concentration equivalent to an optical density at 600 nm of 5, and the samples were heated at 95℃ for 5 min. Proteins were separated by SDS-PAGE and then transferred onto polyvinylidene difluoride (PVDF) membranes. The membranes were incubated with anti-PomA or anti-PomB primary antibodies and the goat anti-rabbit IgG-HRP as a secondary antibody. The protein bands were detected using chemiluminescence procedure with LAS 3000 (Fujifilm).

### Sample preparation and data correction of negative staining images

A peak elution fraction of the PomAB complex separated by size-exclusion chromatography (SEC) was diluted 10-fold in the 20TN100 buffer containing 0.0025%LMNG which is the SEC elution buffer. A 5 μL solution was added to a glow-discharged continuous carbon grid. Excess solution was removed using filter paper, and samples were subsequently stained on the carbon grid with 2% ammonium molybdate. Electron microscopy images were recorded using an H-7650 transmission electron microscope (Hitachi) operated at 80 kV and equipped with a FastScan-F114 CCD camera (TVIPS, Gauting, Germany) at a nominal magnification of 80,000 ×.

## ACKNOWLEDGEMENTS

We thank Dr. Kimika Maki for technical support with electron microscopy. This work was supported in part by JSPS KAKENHI Grant Numbers 23H02447 (to S.K.), and 20H03220 (to M.H.).

